# Structural Covariance of the Duplicated Heschl’s Gyrus: A Sulcal/Gyral Template Morphology Approach

**DOI:** 10.1101/2023.03.29.534799

**Authors:** Mark A. Eckert, Dyslexia Data Consortium

**Author notes:** **Corresponding Author** Mark A. Eckert, Ph.D., Department of Otolaryngology – Head and Neck Surgery, Medical University of South Carolina, Charleston, S.C. 29425, U.S.A.

## Abstract

Heschl’s gyrus (HG) can occur as a single gyrus or with a completely duplicated posterior HG that has been related to a variety of abilities and disorders. Voxel-based studies typically involve the normalization of these qualitatively different HG types, thus making it difficult to evaluate the contribution of sulcal/gyral variability to voxel-based effects and perhaps obscuring some effects. To examine the structural covariance of single and duplicated HG, templates were created for the left single and duplicated HG. Structural covariance analysis with a Jacobian measure of volumetric displacement demonstrated consistent spatial covariance with homologous structure in the right hemisphere across qualitatively different HG morphology. These results suggest that HG duplication is aptly named with respect to cortical structure variation and demonstrate a multi-template approach for studying qualitatively unique brain function and structure linked to perceptual and cognitive functions.

**Highlights:** Qualitatively unique sulcal/gyral features can affect voxel-based analyses.

Heschl’s gyrus is highly variable across people.

Morphology-specific templates were created to study Heschl’s gyrus structural covariance.

Single and duplicated Heschl’s gyrus exhibited a similar pattern of covariance.

## 1. Introduction

Heschl’s gyrus (HG), the site for primary auditory cortex, is highly variable in morphology (Leonard et al., 1998; Penhune et al., 1996). This qualitatively unique variability includes a completely duplicated posterior gyrus that emerges from the circular sulcus immediately behind the anterior HG and runs parallel in an anterolateral direction with the anterior HG. This morphology, which is present in gestation (Pundir et al., 2016), has generated interest because it is more commonly observed among people with right-handedness (Marie et al., 2015), musical training (Benner et al., 2017), dyslexia (Leonard et al., 2001), and schizophrenia (Takahashi et al., 2021). Thus, there are questions about the structural and functional significance of duplicated HG. While there is limited evidence, one study has demonstrated duplicated tonotopic maps across anterior and posterior duplicated HG (Da Costa et al., 2011). Here, we asked if these anterior and posterior HG exhibit the same structural covariance and hypothesized that the anterior and posterior HG exhibit different patterns of structural covariance based on evidence of differences in myelination between the two gyri (Benner et al., 2017). That is, we asked if anatomical variation across the brain is also duplicated across the gyri using a novel sulcal/gyral specific template approach.

The sulcal/gyral specific template method described here was motivated, in part, by the challenges in obtaining quantitative measures of HG duplications. Manual measures are laborious and fraught by limited measurement reliability. Automated measures can be problematic because spatial normalization typically involves normalization to a target with a distinct single HG (e.g., in the MNI template). Motivated by a similar multi-template approach that identified differences in cortical morphology between people with different rhinal sulcus patterns (Xie et al., 2017), morphologically specific single and duplicated HG templates were created in the current study to examine individual differences in cortical morphology, the degree of consistency of structural covariance across different types of HG, and if there were spatial differences in covariance that may provide insight about the role(s) of the duplicated HG, with the overarching goal of demonstrating an approach for dealing with and examining sulcal/gyral variability in neuroimaging studies.

## 2. Material and methods

### 2.1 Participants

This study involved the deformation-based analysis of 194 participants (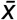 age = 15.51, ± 8.57 years; 35 % female) whose de-identified data were contributed to the Dyslexia Data Consortium. These data were collected in accordance with the Declaration of Helsinki at 14 different research sites and then de-identified prior data sharing, and are thus non-human research data. The primary inclusion criterion was the presence of a clearly visible duplicated left HG that emerged from the medial wall of the Sylvian fissure with an L-shaped Sylvian fissure, which resulted in the identification of 97 cases from a larger set of 1189 images that were examined by the first author who has established expertise characterizing HG morphology (Leonard et al., 2001). Participants with a single left HG were then selected using the R package MatchIt (v4.5.0) to be matched with the duplicated HG cases based on age (*t*_(1,193)_ = 0.16, *p* = 0.87), reported biological sex (*X*^2^ = 0.00, *p* = 1.00), and research site parameters (*X*^2^ = 0.00, *p* = 1.00).

### 2.2. Image Processing

Native space and skull-stripped T1-weighted images were denoised and rigidly aligned to the MNI template using SPM12. Whole brain templates (Supplemental Figure 1) were then created using these images from the single or duplicated HG cases with the ANTS SyN diffeomorphic normalization approach (Avants et al., 2009). This normalization procedure was performed using ten warping steps [Normalization parameters steps 1-10: Cross-correlation metric (mm radius = 4.00); SyN (2, 1, 1); Gaussian regularization (4.00); steps (Step 1: 30 × 90 × 20 × 12; Steps 2-10: 30 × 90 × 30 × 20)]. The initial normalization target was the average of the rigidly aligned images. The average image from each normalization step then served as the target for each subsequent normalization step. Here the goal was to generate template images that reflected the specific morphology of single or duplicated HG. Log Jacobian images were then calculated from the final linear and non-linear warping parameters to place each rigidly aligned images into the single or duplicated HG templates. These images were then smoothed with a Gaussian kernel (full width at half maximum = 8 mm) for statistical analyses, as in our previous work (Eckert et al., 2022).

The average Jacobian determinant was then collected from the unsmoothed Jacobian images from within a region of interest (ROI) for the single (MNI: -34, -30, 17) and duplicated HG (MNI: anterior, -33, -31, 16; posterior, -39, -35, 14) where they first appear in medial plane of section using the SPM toolbox MarsBar. That is, the pial-gray matter boundary of the gyrus was traced along the crown of HG in the most medial section where Heschl’s sulcus was observable and the Jacobian values within that space were averaged for each participant. We also examined results for an ROI that encompassed the entire medial HG at the same sagittal section and observed nearly identical results (Supplemental Figure 2). The same procedure was used for a motor hand bump region that served as a control variable to show spatial specificity of the HG and motor hand bump structural covariance. The pial-gray matter boundary of the hand bump was traced in the most medial position where a complete gyrus was observed to be bounded by sulci (single and duplicated HG template MNI: -34, -27, 51). The Jacobian values represent the extent to which a voxel had to be distorted to fit to the template. Higher Jacobian are observed when voxels need to be reduced in volume to fit to a template. Thus, larger Jacobian values within HG or hand bump regions of interest would be expected to occur for cases with relatively large Heschl’s or hand bump gyri. ANTS SyN normalization was used to warp the single and duplicated HG templates into MNI space and these warps were applied to the respective smoothed and template-specific Jacobian images so that results could be presented in MNI space.

### 2.3. Statistics

This retrospective multi-site study had missing data for behavioral measures, including and a non-word reading measure of phonological decoding (Woodcock-Johnson and Woodcock Reading Mastery Word Attack; 10% missingness) and right-versus non-right-handedness (writing hand and Edinburgh handedness ≤ 0; 10% missingness), which have been associated with duplicated HG (Leonard et al., 2001; Marie et al., 2015). Multiple imputation was used to deal with this missingness, where age, sex, phonological decoding, handedness, and single or duplicated HG type were included in the multiple imputation model. Handedness and phonological decoding associations with HG type were then pooled across 10 imputed datasets.

SPM12 was used to perform structural covariance analyses with the average HG Jacobian determinant information as a predictor of whole-brain voxel-wise Jacobian values. These regression models included research site and the average Jacobian value across the entire brain to control for site effects and globally large or small brain volumes. These types of analyses of course demonstrate nearly perfect correlations within the left HG ROI. To examine the spatial consistency of effects across regions of interest, a *p* < 0.001 threshold was used for each result and thus any region of overlap across analyses had a conjoint *p* value < 1.00e-6. That is, the whole brain structural covariance results had to replicate with a *p* < 0.001 to define significant regions of HG covariance. Similar results were observed when a threshold of p < 0.01 was used (Supplemental Figure 3). To characterize the extent of spatial overlap, the thresholded statistic images were binarized and the corresponding voxel vales were correlated across pairs of images using the corrcoef function in MATLAB (MathWorks, Inc). The R cocor library (v 1.1.4) was used to examine the extent to which the binarized (*p* < 0.001) single HG structural covariance results were significantly more similar to the anterior duplicated HG compared to the posterior duplicated HG.

### 2.4. Research Data

The image processing methods used in this study are relatively standard and have been published previously, including with access to the ANTS code used for normalization (Eckert et al., 2019). The data used in this study are available with completion of an established data use agreement, as required by institutional data sharing procedures.

## 3. Results

### 3.1. Behavioral Associations with HG Morphology

Non-right-handedness was less likely to occur (*X*^2^ = 4.49, *p* = 0.039) with a duplicated HG (4 %) than a single HG (13 %). Lower phonological decoding was also observed (*F*_(1,193)_ = 4.95, *p* = 0.027) with a duplicated HG (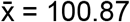, ± 15.90) compared to a single HG (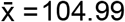, ± 15.47). That is, small effect handedness and phonological decoding associations were observed with qualitative HG sulcal/gyral variability. There was no evidence that the patterns of structural covariance described next could significantly explain within HG group variation in handedness or phonological decoding.

### 3.2. HG Structural Covariance

Figure 1A shows that Jacobian volume of the single, duplicated anterior, and duplicated posterior HG exhibited significant correlations with Jacobian volume of the right HG (*p* < 0.001 for each analysis). That is, a larger left HG occurred with a larger right HG regardless of the type of left HG. There were no significant differences in strength of left HG correlation with the right HG between single and duplicated HG (*p*s > .512 for differences between *r*s = 0.416, 0.426, 0.492, respectively). A similar association with homologous contralateral structure is shown in Figure 1B for the motor hand bump where a larger left hemisphere hand bump occurred with a larger right hemisphere hand bump.

**Figure 1.**
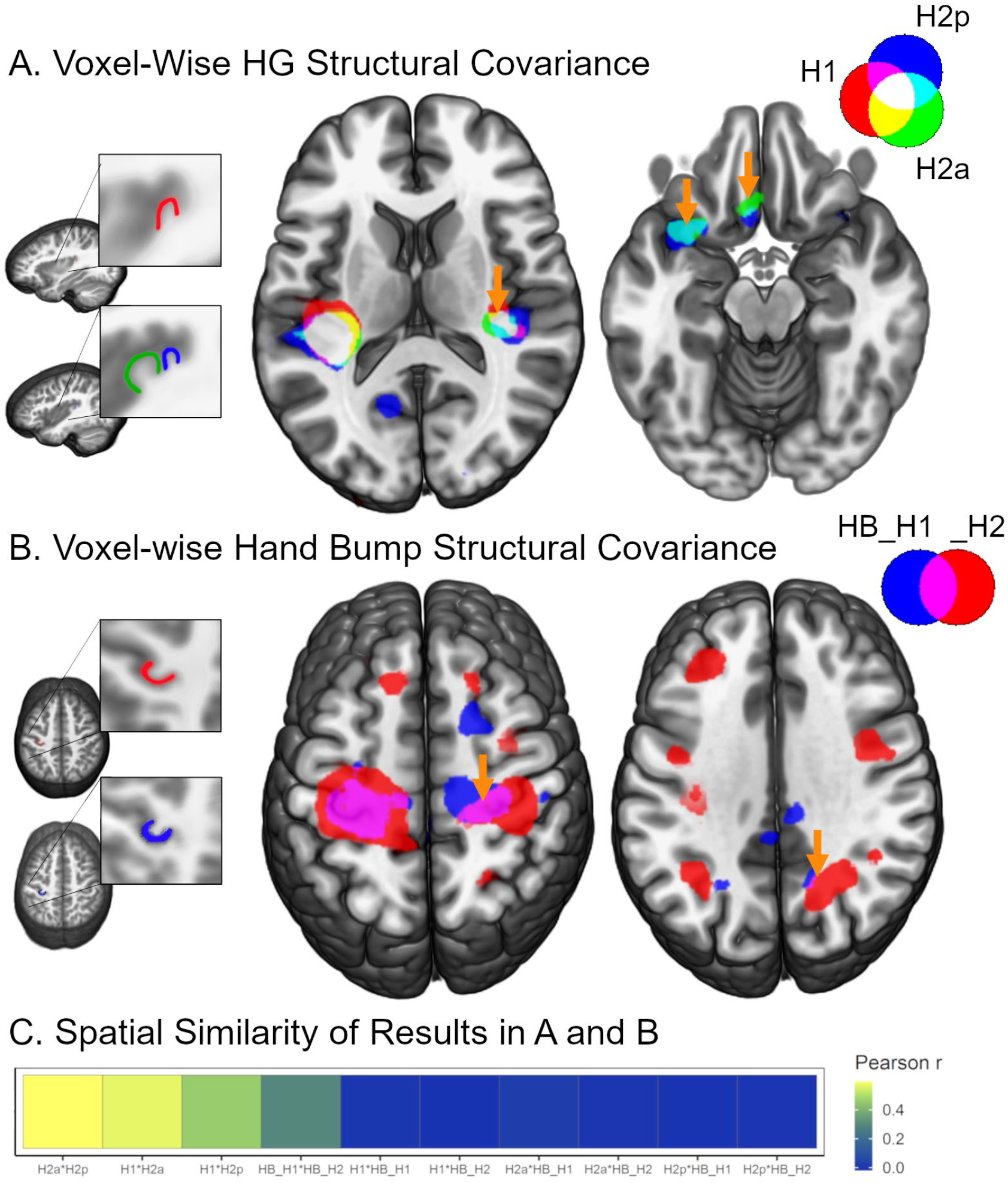
Spatial similarity of the voxel-wise associations for each ROI pair. A. Overlap of the structural covariance results (arrows) for the left single HG cases (red), as well as the anterior (green) and posterior (red) duplicated HG cases, including for right auditory cortex. B. Similar overlapping contralateral effects (purple) were observed for the left motor hand bump across single HG (red) and left duplicated HG (blue) cases. C. The results images shown in A and B were correlated to characterize the extent of spatial similarity for each pair of results, where relatively a large association was observed for the anterior and posterior HG for cases with a duplicated HG. H1: single HG; H2a: anterior duplicated HG; H2b: posterior duplicated HG; HB_H1: Hand bump in the single HG cases; HB_H2: Hand bump in the duplicated HG cases; Results images were thresholded at *p* < 0.001. The green HG outline in A is presented for color reference to the structural covariance results but this ROI was more medial where it first appears emerges medially.

Jacobian volume in the anterior and posterior duplicated HG also exhibited spatially overlapping associations with Jacobian volume in the left frontoinsular and orbitofrontal regions. For example, the average *r* values within the overlapping frontoinsular region in the anterior (*r* = 0.421, *p* < 0.001) and posterior (*r* = 0.451, *p* < 0.001) duplicated HG were significantly greater than for the single HG (*r* = -0.071, *ns*) using a Fisher’s Z comparison (*p*s < 0.0003). Aside from the right HG, there were no other brain regions where the single HG and duplicated HG exhibited significant spatially corresponding associations. This pattern of spatially overlapping and non-overlapping results across regions of interest is consistent with the strength of associations between pairs of ROI results (Figure 1), where there was a significant association between the duplicated HG result maps (*r* = 0.595, *p* < 0.001). The single HG structural covariance result map was significantly related to the map for the anterior single HG (*r* = 0.567, *p* < 0.001), and to a lesser extent with the posterior duplicated HG result map (*r* = 0.461, *p* < 0.001). Comparison of these associations demonstrated a significantly stronger association between the single HG and anterior duplicated HG than the single HG and posterior duplicated HG (*z* = 273.95, *p* < 0.001). Figure 1A suggests that one reason for this difference is the posterior duplication association with visual cortex. This visual cortex result was not significant after Bonferroni correction. To put the difference in HG spatial covariance results in context, the structural covariance result maps for the hand bump control region exhibited substantively less similarity (*r* = 0.294, *p* < 0.001) than the HG structural covariance results maps.

## 4. Discussion

A sulcal/gyral specific template approach demonstrated that HG morphology exhibits a common pattern of structural covariance across single, anterior duplicated, and posterior duplicated morphological features. There was more spatial consistency in the pattern of structural covariance for the single and anterior duplicated HG compared to the single and posterior duplicated HG, but this difference was smaller than the difference in magnitude of spatial correlation for these associations relative to the hand bump spatial covariance associations between templates. There were, however, unique patterns of association within frontoinsular and orbitofrontal cortex for the duplicated compared to the single HG cases. Together, the results provide additional evidence that duplicated anatomical HG may have common developmental origins and demonstrate a multi-template framework for examining brain structure and function.

### 4.1. Heschl’s Gyrus Homology

A putative estimate of myelin (T1/T2 ratio) was lower in participants with a posterior duplication compared to their anterior duplication (Tzourio-Mazoyer et al., 2019). This observation motivated the hypothesis that anterior and posterior HG duplications would exhibit differences in structural covariance, perhaps due to differences in the patterns of structural connectivity occurring with differences in myelinated fibers terminating/originating in HG. However, the spatial pattern of structural covariance was highly similar for the two types of HG morphology. Moreover, the patterns of association in duplicated HG were also similar to the patterns observed in the cases with a single HG. It may be that HG T1/T2 ratio covariance with other brain regions yields similar results to those described here.

### 4.2. Spatial Specificity of Structural Covariance

HG and the motor hand bump Jacobian data exhibited significant associations with the contralateral homologous structure. These effects were remarkable in their spatial specificity relative to other brain regions and were not dependent on whole-brain Jacobian values, which were largely correlated with voxel values at the cortical boundary. These results are generally consistent with functional connectivity imaging studies demonstrating a relatively high degree of functional covariance between homologous structures between hemispheres (Andoh et al., 2015). The duplicated HG did exhibit frontoinsular and orbitofrontal associations that were not observed in the single HG participants, and while not clearly related to phonological decoding, these are regions that have been consistently related to reading disability (Eckert et al., 2016).

### 4.3. Behavioral Significance

The single HG cases had a greater frequency of non-right-handed cases and people with better phonological decoding compared to the duplicated HG cases. Previous studies have selected for handedness or reading disability groups and then examined differences in HG morphology. Here, selection was based on the presence of a clearly observable duplicated HG and the research site, age, and sex matching was performed to identify the single HG cases. For this reason, it may be remarkable even small effects with handedness and phonological decoding were observed. However, there was limited evidence that individual variation in the Jacobian data was related to these variables within HG groups. Moreover, the non-specific HG duplication associations such as reading disability and schizophrenia suggest that HG duplications are a marker of non-canonical development rather than a specific causal influence for disability.

### 4.4. Limitations

One limitation of the sulcal/gyral specific template approach, particularly with Jacobian data, is that the values are dependent on the other participants that contribute to each template, thus making it difficult to compare participants across templates. For this reason, it was not possible to clearly compare Jacobian values between HG groups. Deformation-based analysis using data from a template composed of all participants was considered but it was not clear that this approach would be additive given the consistency of results across templates. A better approach for future studies may be to use the multi-template method and apply the normalization warped to independently derived metrics such as T1/T2 ratio.

This study of retrospective data was also limited in the availability of consistent supporting demographic and behavioral data across the cases. While multiple imputation was used to deal with missingness, there was inconsistency in data collection across sites. For example, writing hand and quantitative hand preference data were used to define handedness, so caution is appropriate when interpreting the handedness results.

### 4.5. Future Applications

The current study was designed to use a sulcal/gyral specific template approach to advance understanding about HG morphology. The same approach can be used for other cortical regions that exhibit individual variability in sulcal/gyral morphology (e.g., for people with and without a paracingulate sulcus) to examine the structure and function of these regions. Functional imaging studies of auditory cortex, for example, often include analysis of responses to auditory stimuli in a common coordinate space without considering differences in HG morphology. The approach described here may enhance sensitivity for group analyses. Moreover, a template matching approach could be used to identify the template that provides the best fit for a given morphology so that expertise in phenotyping anatomical features is not critical for this type of analysis or to phenotype HG morphology to study why duplicated HG occur with a variety of disorders, handedness, and musical expertise.

## 5. Conclusions

A HG morphology template approach was used to establish a high degree of similarity in structural covariance across people with qualitatively different HG morphology. The results, particularly the spatial covariance with the contralateral auditory cortex, suggest that HG duplication is an apt term for the completely separate gyrus that appears behind the primary HG. The template approach used here can also be used to examine the morphology of other sulcal/gyral features, as well as consider the functional significance of differences sulcal/gyral morphology that have been a source of intrigue because of associations with normative and atypical behavior.

## Acknowledgements

This work was supported by HD069374. This investigation was conducted in a facility constructed with support from Research Facilities Improvement Program (C06 RR014516) from the National Center for Research Resources, National Institutes of Health.

## Notes

### Competing Interest Statement

The authors have declared no competing interest.

