## Supplemental Materials for "Structural Covariance of the Duplicated Heschl’s Gyrus: A Sulcal/Gyral Template Morphology Approach"

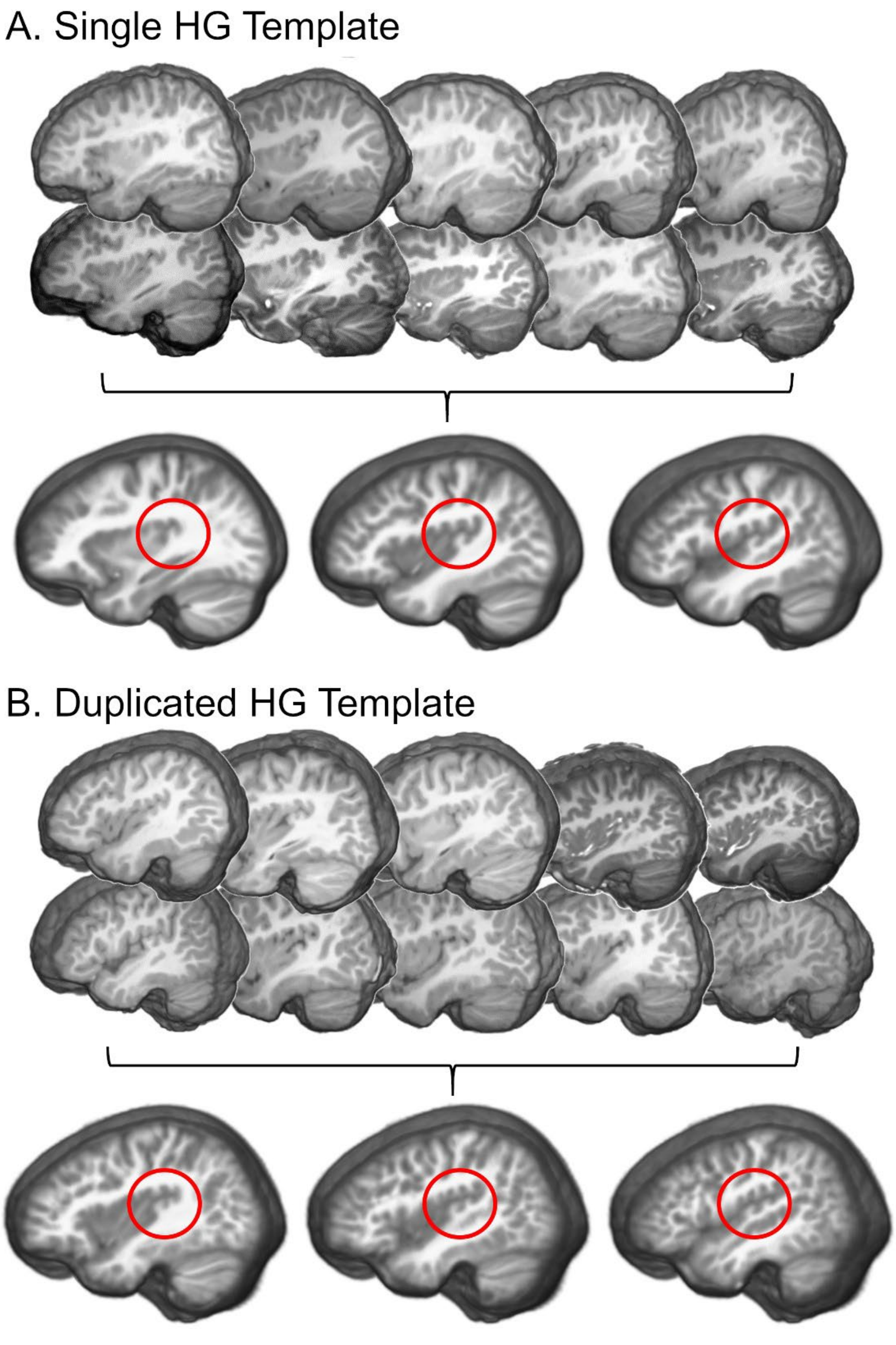


**Supplemental Figure 1.** Single and Duplicated Heschl's gyrus (HG) templates. Images from 97 participants with a single HG (A) and 97 with a duplicated HG (B) who were age, sex, and research site matched were used to generate HG morphology specific templates using ANTS SyN normalization. Red circles highlight HG. Example cases for each HG template are shown to demonstrate the single and double HG features contributing to each template where the red circle highlights HG.

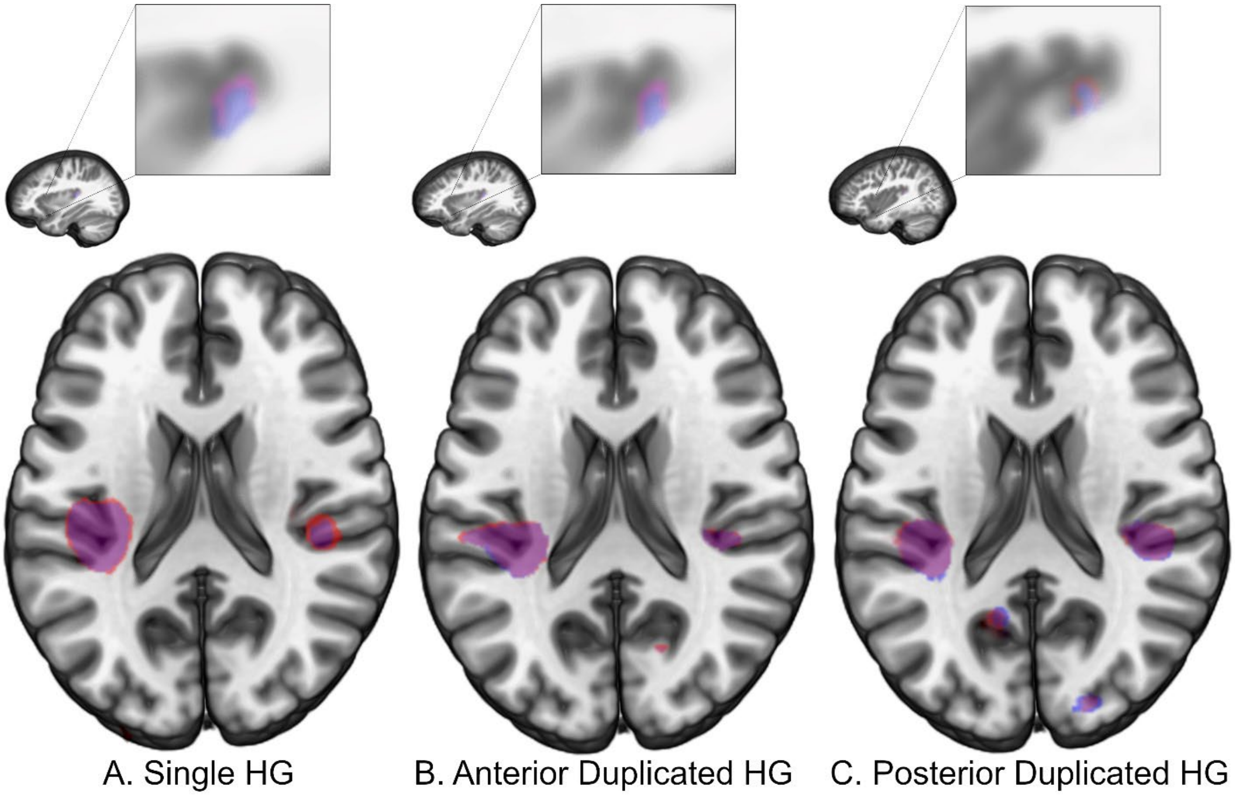


**Supplemental Figure 2**. Consistent spatial covariance results across HG regions of interest and across HG morphology (A-C). The insets at top show the regions of interest where either the boundary of Heschl's gyrus was defined as the region of interest or the entire cross-section of the Heschl's gyrus was defined as the region of interest. Structural covariance results are shown for each region of interest and demonstrate a high degree of overlap (purple) across region of interest analyses for each type of Heschl's gyrus.

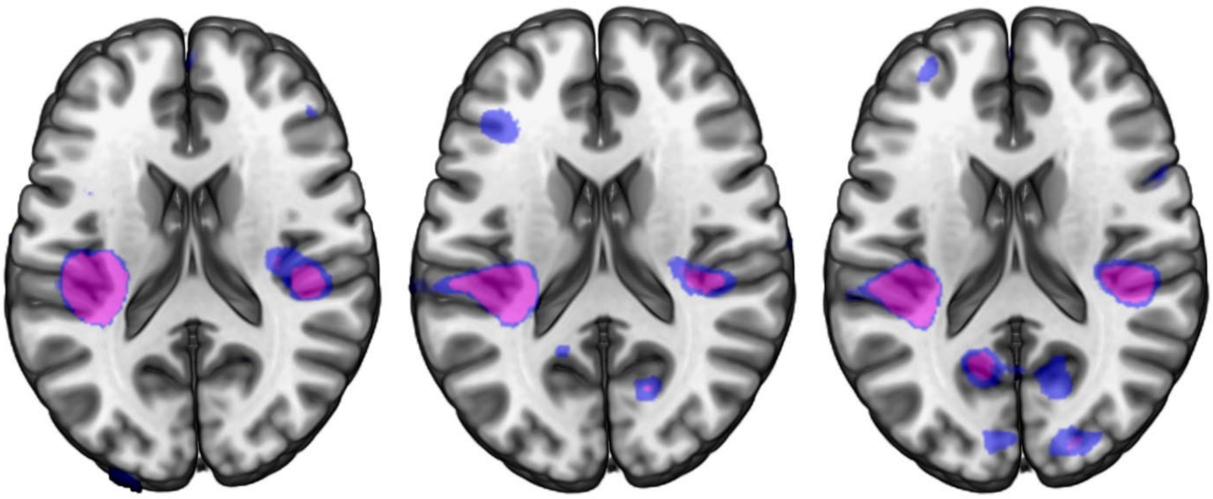


**Supplemental Figure 3**. The same pattern of spatial covariance results was observed across HG regions of interest for different statistical thresholds (red: $p < 0.001$; blue: $p < 0.01$; purple: overlap or red and blue). Intraparietal sulcus effects overlap for the anterior and posterior duplicated HG but caution should be used interpreting these more modest effects.
